# *De novo* transcriptome analysis of dermal tissue from the rough-skinned newt, *Taricha granulosa*, enables investigation of tetrodotoxin expression

**DOI:** 10.1101/653238

**Authors:** Haley C. Glass, Amanda D. Melin, Steven M. Vamosi

## Abstract

**Background:** Tetrodotoxin (TTX) is a potent neurotoxin used in anti-predator defense by several aquatic species, including the rough-skinned newt, *Taricha granulosa*. While several possible biological sources of newt TTX have been investigated, mounting evidence suggests a genetic, endogenous origin. We present here a *de novo* transcriptome assembly and annotation of dorsal skin samples from the tetrodotoxin-bearing species *T. granulosa*, to facilitate the study of putative genetic mechanisms of TTX expression.

**Findings:** Approximately 211 million read pairs were assembled into 245,734 transcripts using the Trinity *de novo* assembly method. Of the assembled transcripts, we were able to annotate 34% by comparing them to databases of sequences with known functions, suggesting that many transcripts are unique to the rough-skinned newt. Our assembly has near-complete sequence information for an estimated 83% of genes based on Benchmarking Universal Single Copy Orthologs. We also utilized other comparative methods to assess the quality of our assembly. The *T. granulosa* assembly was compared with that of the Japanese fire-belly newt, *Cynops pyrrhogaster*, and they were found to share a total of 30,556 orthologous sequences (12.9% gene set).

**Conclusions:** We provide a reference assembly for *Taricha granulosa* that will enable downstream differential expression and comparative transcriptomics analyses. This publicly available transcriptome assembly and annotation dataset will facilitate the investigation of a wide range of questions concerning amphibian adaptive radiation, and the elucidation of mechanisms of tetrodotoxin defense in *Taricha granulosa* and other TTX-bearing species.

## Data Description

### Context

In recent years, there has been rapid progress in massive parallel sequencing technologies, but there remain many challenges for genomic studies of non-model organisms [1, 2]. It is especially difficult to study the genomics of species that have highly repetitive genomes, particularly amphibians, and it can be insurmountable to produce genome assemblies due to computing resource limitations [3]. To date there are only seven amphibian genomes published in the NCBI database, compared to e.g., 654 mammal and 600 insect genomes [4]. A solution to this problem is to utilize a transcriptomics or RNA-sequencing (RNA-seq) approach from high-throughput sequencing data build from high-quality RNA. One goal of RNA-seq is to interpret the functional elements of the genome, which makes this a feasible solution for studying genes from species with highly repetitive genomes [5]. Moreover, this approach allows the *de novo* assembly and analysis of the transcriptome without mapping to a reference genome [6]. *De novo* transcriptome assembly methods can outperform genome-guided assemblies in non-model organisms and are especially useful for diverged species or those with complex genomes [7].

Researchers would benefit from having improved genetic and genomic resources for one amphibian species that has been the subject of much historical and contemporary work and is now a textbook story of predator-prey interaction: the rough-skinned newt, *Taricha granulosa*. This urodelian amphibian, endemic to the west coast of North America, possesses a potent neurotoxin known as tetrodotoxin (TTX) that is primarily excreted through their dorsal skin as a predator deterrent [8]. There is significant variation in TTX concentration among populations of newts, and researchers had previously suggested this is attributed to predation pressure from garter snakes, *Thamnophis sirtalis* [i.e., 9, 10, 11]. However, new work has suggested that TTX variation in rough-skinned newts may instead be due to neutral population divergence, whereas snake resistance is an evolutionary response to prey toxicity [12]. This defense mechanism has also convergently evolved in several other diverse species including puffer fish, blue ringed octopus, and moon snails [13].

Although the toxicity of rough-skinned newts has been well studied over the past few decades, the source of toxin production has been widely debated. First, it was suggested that TTX is bio-accumulated as a result of diet, but this was rejected when newts that were kept in captivity and fed a TTX-free diet retained their toxin concentrations over time [14]. Next, Lehman et al. [15] rejected the hypothesis that bacterial endosymbionts are responsible for the observed toxicity after discovering that none of the bacteria present on newt skin were capable of producing TTX. These findings led to the hypothesis that tetrodotoxin biosynthesis is under genetic control and toxins are synthesized within the individual. However, a satisfactory determination of the genetic mechanisms responsible for TTX expression remains lacking, despite the fact that previous studies suggest an endogenous origin [16].

In addition to the lack of information regarding the origin of tetrodotoxin in rough-skinned newts, they also have very limited genetic data published in general. We are not aware of any work published on high-throughput sequencing data from this species, and previous studies have focused only on mitochondrial markers, allozymes, and microsatellites [i.e., 17, 18, 19, 20]. While rough-skinned newts are presently listed as a species of Least Concern, a subspecies known as the Mazama newt (*Taricha granulosa mazamae*) is currently threatened by invasive crayfish in Crater Lake National Park in Oregon, USA [21]. Due to the sensitivity of amphibians to invasive species such as crayfish, as well as other anthropogenic disturbances, it will be crucial to have more robust genetic resources of rough-skinned newts available.

Our goal is to provide a high-quality transcriptome from the dorsal tissue of a newt species known to produce high quantities of tetrodotoxin, in order to make a crucial step towards elucidating the genetic mechanisms of TTX expression. Here, we present the transcriptome assembly and annotation of *Taricha granulosa* using Illumina RNA-sequencing technology and bioinformatics tools. Genetic and genomic references for rough-skinned newts are sparse and, to our knowledge, this is the first report on transcriptome analyses for the species. This work provides a valuable resource for studying the chemical ecology of tetrodotoxin in *T. granulosa* as well as for other TTX-bearing species. In addition, this will expand our understanding of salamander genes in general.

### Sample Collection and Preparation

Four male rough-skinned newts were collected from different lakes in British Columbia, Canada either by net or in a minnow trap. Following a non-lethal sampling method [22], a 2mm skin biopsy tool (Robbins Instruments, USA) was used to remove a skin sample from the posterior dorsolateral area of each individual, and the newts were returned to their original habitat. The skin sample was immediately placed in a tube of RNAlater solution (Thermo Fisher Scientific, USA), which was kept at room temperature for approximately 24 hours. The tubes were then placed in a liquid nitrogen dewar at approximately −190C before transportation back to the University of Calgary for storage at −80C.

Total RNA was isolated and purified with the RNeasy micro extraction kit (Qiagen, USA) using a modified protocol in which the tissue samples were macerated with a glass tissue grinder without freezing in liquid nitrogen. Individual libraries were constructed using the TruSeq Stranded mRNA Library Prep kit (Illumina, USA). The libraries were sequenced (100-bp paired-end) on an Illumina HiSeq 4000 at a sequencing depth of approximately 51-62M reads per library. Library preparation and sequencing was performed at the Genome Quebec Innovation Centre. Raw sequencing reads can be found at the National Center for Biotechnology Information (NCBI) Short Read Archive (SRA) under BioProject Accession PRJNA505885.

### *De Novo* Transcriptome Assembly

Raw reads from the four individuals were first pooled into a single fasta file, representing a combined total of 230,544,185 reads. To remove low quality reads, quality control was performed with Trimmomatic v0.36 [23] using the following parameters: (i) Trimming of bases at the leading and trailing ends of sequences with a phred+33 quality score below 20, (ii) a 4-base sliding window scan to remove read fragments with an average quality per base below 20, and (iii) removal of reads below 36 base pairs long. After trimming, 91.47% of read pairs (N=210,868,103) remained, and only reads with both pairs remaining were used for the assembly. The quality-filtered reads were then used to perform a *de novo* transcriptome assembly with Trinity v2.6.5 [6, 24] using the default parameters. The reads were assembled into 245,734 transcripts and represent an average length of 837.57 bp, median length of 339, N50 length of 2093, and GC content of 44.96% (Table 1, Fig. S1: Additional File 1).

**Table 1.** Assembly statistics for the *de novo* assembled transcriptome of the rough-skinned newt.

| **Assembly Statistics** |  |
| --- | --- |
| Number of Filtered Read Pairs | 210,868,103 |
| Assembly Size, bp | 205,818,378 |
| Number of Trinity Transcripts | 245,734 |
| Number of Trinity Genes | 167,028 |
| GC (%) | 44.96 |
| Contig N10 | 6,738 |
| Contig N20 | 4,914 |
| Contig N30 | 3,799 |
| Contig N40 | 2,887 |
| Contig N50 | 2,093 |
| Median Contig Length, bp | 338 |
| Average Contig Length, bp | 837.57 |

### Assembly Assessment

We utilized several methods to assess the quality of the *de novo* assembly we generated. The quality filtered reads were aligned to our assembly with Bowtie2 [25], revealing that approximately 98% of paired reads were properly mapped. Next, we estimated the assembly completeness using BUSCO v3.2.0 [26] by determining the conserved ortholog content with the Tetrapod OrthoDB database [27]. Benchmarking Universal Single-Copy Orthologs (BUSCOs) were identified which revealed that our transcriptome has near-complete genes for 82.5% of the *T. granulosa* genome [46.3% complete and single-copy (N=1830), 36.2% complete and duplicated (N=1428)], with 5.7% fragmented (N=225) and 11.8% missing (N=467) (Fig. S2: Additional File 1). The duplication seen in our BUSCO results is also similar to that of *Cynops pyrrhogaster* [28] and *Ambystoma mexicanum* [29], and could be due to several reasons. First, amphibians are known to have complex, highly duplicated genomes [30]. The Trinity assembly may also include isoforms of the same gene or allelic heterozygote transcripts from different individuals.

We also measured the similarity between our assembly and that of retinal tissue from the Japanese fire belly newt, *C. pyrrhogaster* [28], since these species are both from the Salamandridae family and had comparable assembly statistics. Orthologs between the rough-skinned newt and eastern newt transcriptome assemblies were identified using a reciprocal best-hit BLAST approach. A total of 30,556 orthologous sequences between the two transcriptomes were detected with a query coverage above 70%, which represented approximately 12.9% of *C. pyrrhogaster* transcripts. The number of orthologous sequences is higher than in other salamander species reciprocal best-hit blast comparisons [31] and could be because they both belong to the Salamandridae family and share a more recent common ancestor than some of the other cross-family orthology comparisons.

Additionally, we compared the assembly statistics from our transcriptome to those of six other species to benchmark our assembly completeness (Table 2). These studies were selected for comparison in order to represent a variety of urodelian amphibian species with completed transcriptome assemblies. The *de novo* transcriptome assembly of *T. granulosa* yielded comparable assembly and annotation statistics to other salamander assemblies. Specifically, it had similar BUSCO results to *C. phyrrogaster* and *A. mexicanum*.

**Table 2.** Assembly information and BUSCOs of this study and other studies of closely related species.

| Species | Assembled Bases, bp | Number of Transcripts | Number of Genes | BUSCOs |
| --- | --- | --- | --- | --- |
| *Taricha granulosa*,<br>Rough-skinned newt | 205,818,378 | 245,734 | 167,028 | C: 82.5% [D: 36.2%], F: 5.7%, M: 11.8% |
| *Cynops pyrrhogaster*,<br>Japanese fire belly newt<sup>a</sup> | Not reported | 237,120 | Not reported | C: 82% [D: 34%], F: 4.6%, M: 13% |
| *Nopthothalamus viridescens*, Eastern newt<sup>b</sup> | Not reported | 120,922 | Not reported | C: 30% [D: 7.0%], F: 10%, M: 58% |
| *Lissotriton boscai*, Bosca's newt<sup>c</sup> | 173,736,688 | 153,270 | Not reported | Not Reported |
| *Ambystoma mexicanum*,<br>Mexican axolotl<sup>d</sup> | Not reported | 1,554,055 | 1,388,798 | C: 88% [D: 53%],<br>F: 4.5%, M: 7.3% |
| *Andrias davidianus*,<br>Chinese giant<br>salamander<sup>e</sup> | 128,175,999 | 158,103 | 132,912 | Not Reported |
| *Bolitoglossa ramosi*,<br>Ramos' mushroomtongue<br>salamander<sup>f</sup> | 654,673,506 | 577,037 | 433,809 | C: 78.2% [D:<br>2.4%], F: 9.9%,<br>M: 11.9% |

<sup>a</sup>[28];<sup>b</sup>[37];<sup>c</sup>[38];<sup>d</sup>[29];<sup>e</sup>[39];<sup>f</sup>[31]. Statistics from the de novo assembly and BUSCOs (C: Complete, D: Duplicated, F: Fragmented, M: Missing) of our *T. granulosa* transcriptome together with transcriptome assemblies of related species. Complete BUSCOs include both complete & single-copy and complete & duplicate sequences.

### Functional Annotation and Gene Ontology

To better understand putative functions of assembled transcripts, we conducted functional annotation and assigned Gene Ontology (GO) terms with the Trinotate v3.1.1 pipeline [29]. TransDecoder v5.5.0 was first used to locate candidate coding regions and identify those with the longest open reading frame [32]. Contigs were characterized via BLASTx and BLASTp sequence homology searches against UniProt/SwissProt NCBI NR protein database, using BLAST+ v2.4.0 with an e-value cutoff of e–5 [33]. We based our choice of e-value following methods for the annotation of the axolotl transcriptome by the authors of the Trinotate pipeline [29]. Additionally, PFAM functional domains were identified with HMMER v3.0 [34], transmembrane domains with TMHMM v2.0 [35], and signal peptides with SignalP v4.1 [36]. Gene Ontology terms were also assigned via Trinotate v3.1.1 and were based on best matches in the UniProt/SwissProt database. The results from these analyses were loaded into an Sqlite database to generate a Trinotate functional annotation report.

There were a total of 82,960 annotated transcripts for *T. granulosa*, representing about 33.8% of the 245,734 *de novo* assembled transcripts. This annotation rate is similar to the 36.7% of 237,120 transcripts annotated for *C. phyrroghaster* [28]. Unannotated transcripts could indicate artifacts of misassemblies or non-coding RNAs, but many may also be novel genes with unknown function due to the general lack of knowledge about newt proteins. A total of 48,309 transcripts were assigned at least one GO term, which covered various functional pathways (Table 3). Of those, approximately 48.6% were classified as biological processes, 23.0% as molecular functions, and 28.4% as cellular components. Many GO terms could be assigned to one transcript, so there was significant overlap between the three categories (Fig. S3: Additional File 1).

**Table 3.**
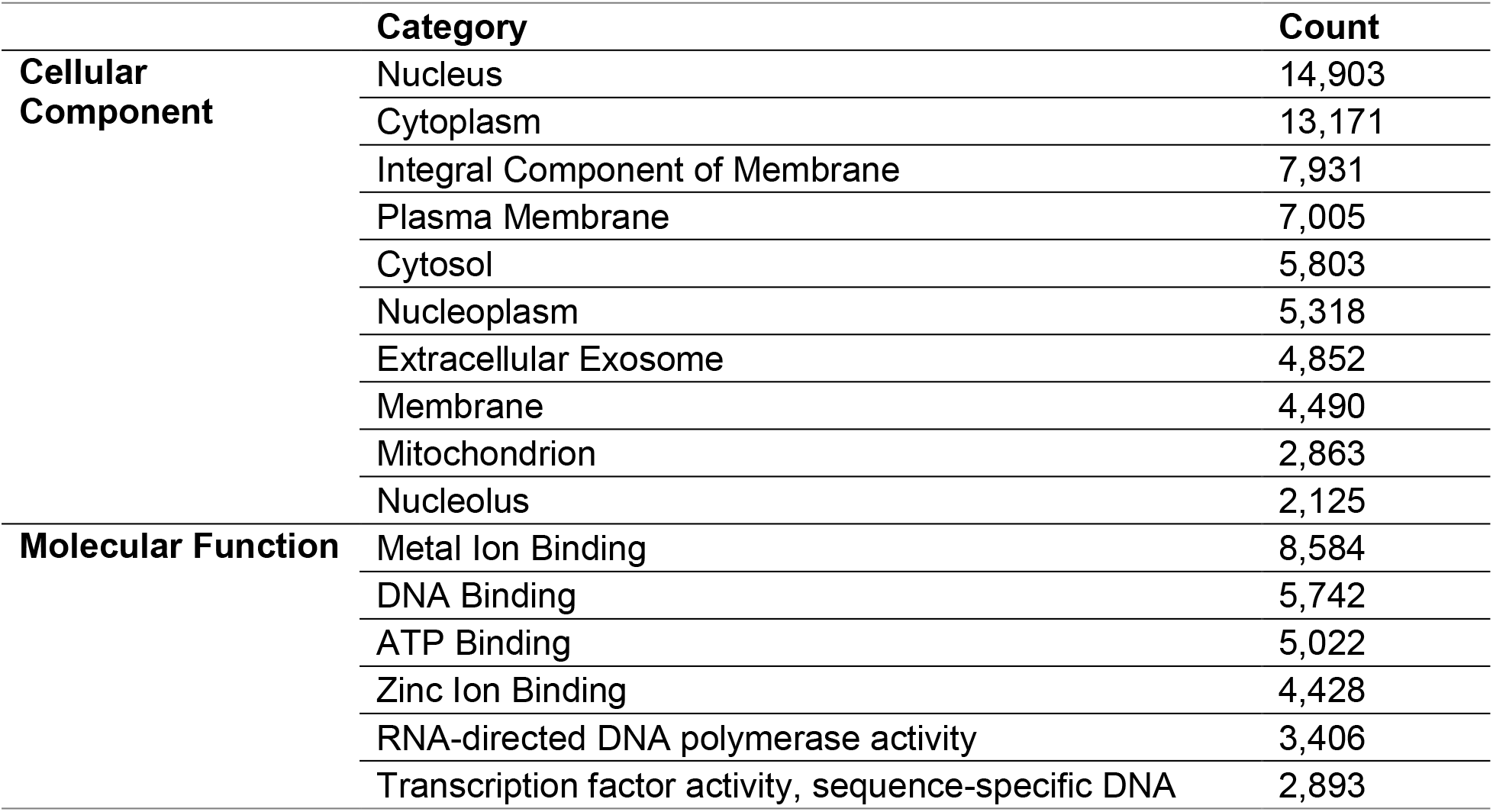
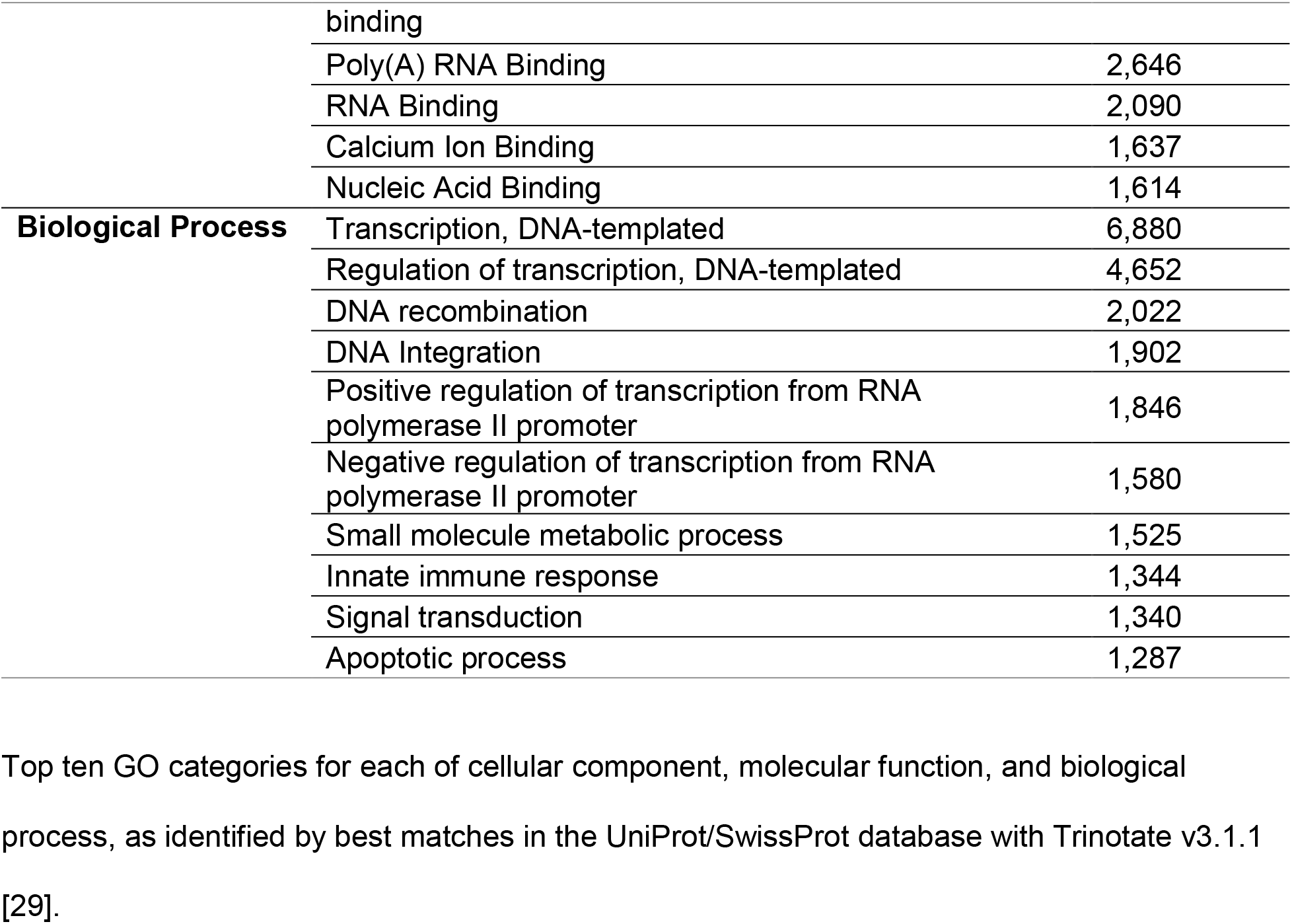
Top gene ontology categories.

## Conclusions

To our knowledge, we present the first report of a transcriptome assembly and annotation of dorsal skin from the tetrodotoxin-bearing rough-skinned newt, *Taricha granulosa*. Rough-skinned newts are known to excrete high concentrations of a potent neurotoxin called tetrodotoxin, but previous studies have been unable to identify an endogenous origin in this species. The transcriptome data provided here will be a valuable resource in elucidating potential genetic mechanisms of tetrodotoxin expression in *T. granulosa* and other TTX-bearing species. Additionally, our work may be useful to other researchers studying this species for other purposes such as population genetics analyses or comparative transcriptomics. This data will also advance the availability of amphibian genomic resources, and enable researchers to continue expanding our knowledge of amphibian genes.

## Availability of Supporting Data

The raw sequencing reads are available in the NCBI Short Read Archive (SRA) under BioProject Accession PRJNA505885. This Transcriptome Shotgun Assembly project has been deposited at DDBJ/EMBL/GenBank under the accession GHKF00000000. The version described in this paper is the first version, GHKF01000000.

### Additional Files

Additional File 1: Figure S1: Histogram showing the distribution of transcripts assembled by Trinity v2.6.5, binned by length.

Additional File 1: Figure S2: Benchmarking Universal Single-Copy Orthologs (BUSCOs) identified by comparing the assembly to the Tetrapod OrthoDB database. The BUSCOs were 46.4% complete and single-copy (N=1834), 36.6% complete and duplicated (N=1445), 5.5% fragmented (N=218), and 11.5% missing (N=453).

Additional File 1: Figure S3: Venn diagram of Gene Ontology (GO) categories of the annotated transcripts, assigned using the Trinotate pipeline. The values represent the number of transcripts with a GO assignment from each of 3 categories: Biological Process, Molecular Function, and Cellular Component.

## Declarations

## Abbreviations

TTX: Tetrodotoxin
RNA-seq: RNA sequencing
bp: base pairs
BUSCO: Benchmarking Universal Single Copy Orthologs
GO: Gene Ontology
NCBI: National Center for Biotechnology Information
SRA: Short Read Archive
TSA: Transcriptome Shotgun Assembly

## Ethics Statement

Ethics approval was provided by the University of Calgary under Animal Use Protocol AC15-0033 and sample collection was performed in accordance with BC Wildlife Act Permit NA17-263401.

## Competing Interests

We declare no competing interests.

## Author Contributions

HG completed field work and collected samples, isolated RNA and prepared samples for sequencing, performed transcriptome assembly and analysis. HG, AM, and SMV participated in study design, manuscript draft preparation, and review and approval of the manuscript for final submission.

## Acknowledgments

This work was funded by a Natural Sciences and Engineering Research Council of Canada grant (03915-2018) to AM, infrastructure provided by the Canada Research Chairs program (231257) and Canadian Foundation for Innovation (35776) to AM, and internal research funds provided by the University of Calgary to SMV. This research was also enabled in part by access to computing resources provided by WestGrid [40] and Compute Canada [41], along with the Helix cluster from the University of Calgary’s Cumming School of Medicine. We would also like to thank Jade Spruyt for her assistance in the field and the lab and Gwen Duytschaever for her ongoing support in the lab.

