## Additional File 1 for "*De novo* transcriptome analysis of dermal tissue from the rough-skinned newt, *Taricha granulosa*, enables investigation of tetrodotoxin expression"

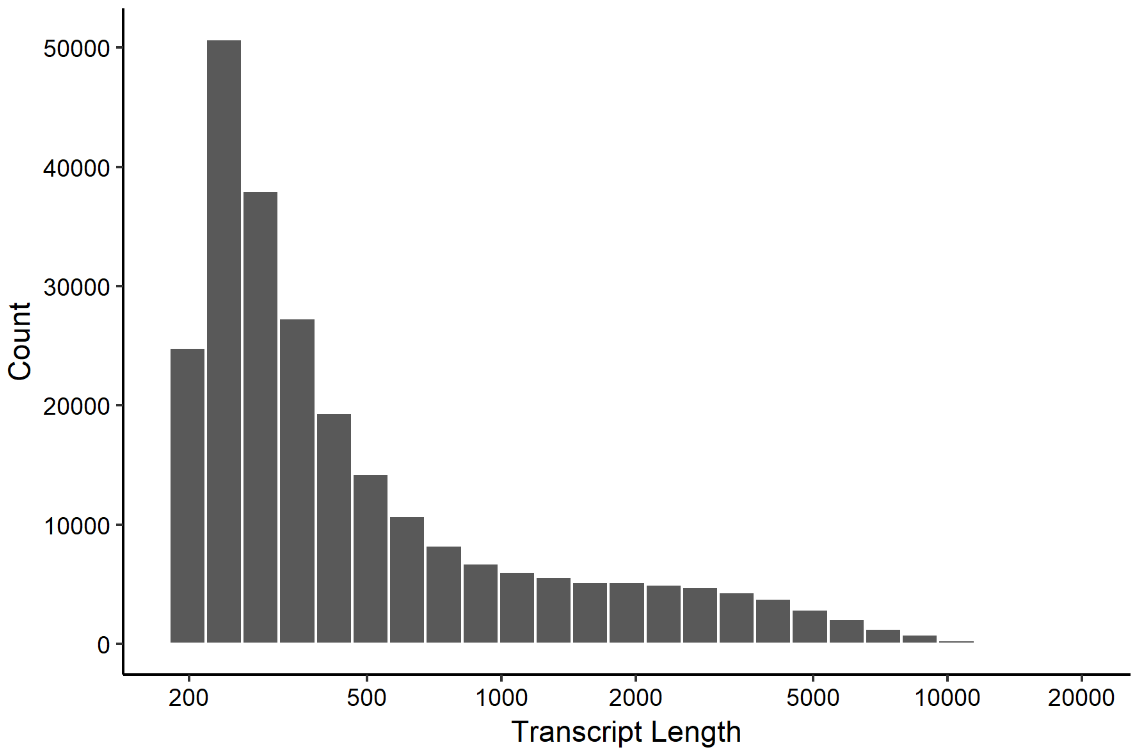


**Figure S1.** Histogram showing the distribution of transcripts assembled by Trinity v2.6.5 [1, 2] binned by length.

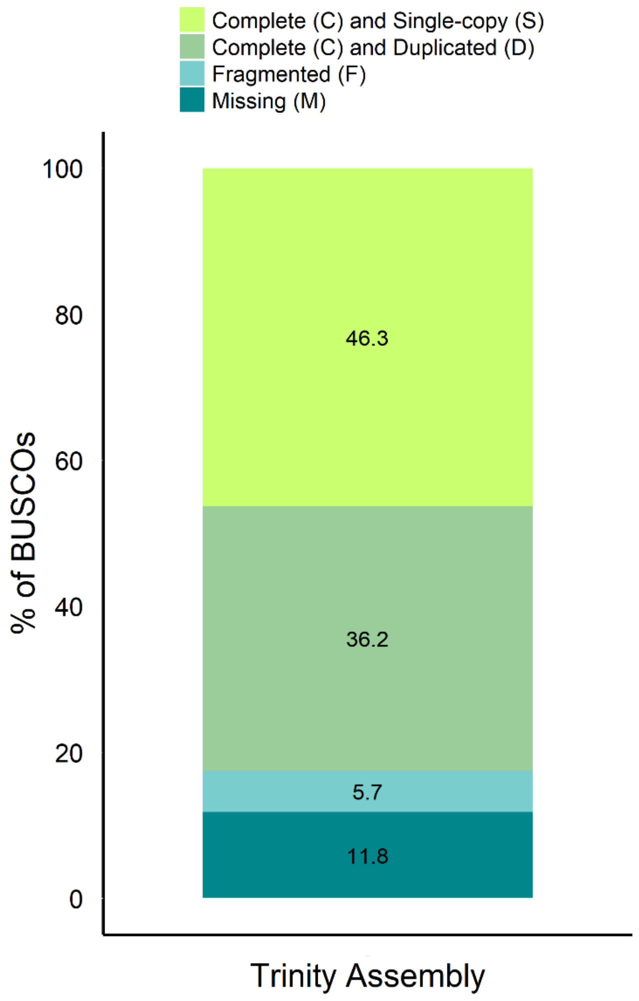


**Figure S2.** Benchmarking Universal Single-Copy Orthologs (BUSCOs) identified by comparing the assembly to the Tetrapod OrthoDB database [3, 4]. The BUSCOs were 46.3% complete and single-copy (N=1830), 36.2% complete and duplicated (N=1428), 5.7% fragmented (N=225), and 11.8% missing (N=467).

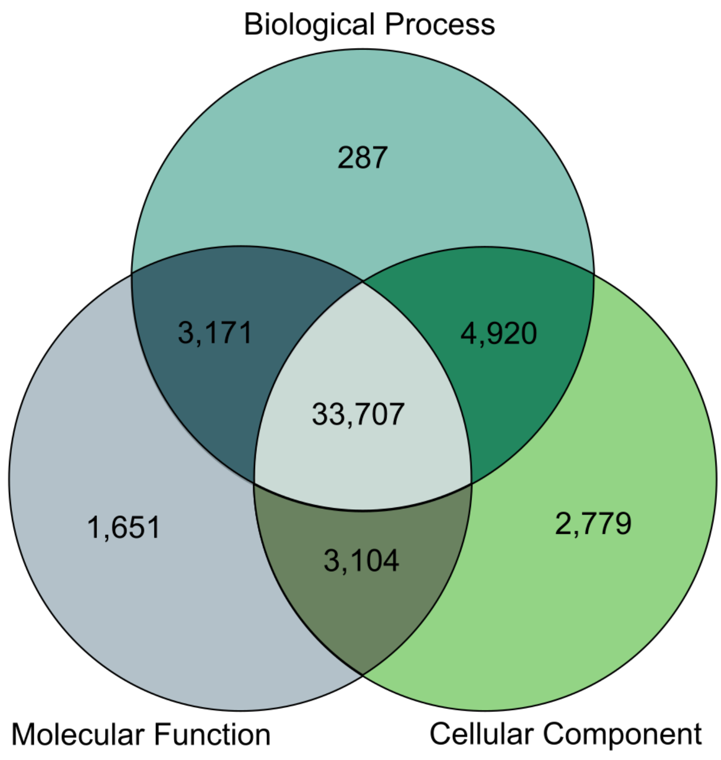


**Figure S3.** Venn diagram of Gene Ontology (GO) categories of the annotated transcripts, assigned using the Trinotate pipeline [5]. The values represent the number of transcripts with a GO assignment from each of 3 categories: Biological Process, Molecular Function, and Cellular Component.
